# Potent Efficacy of Computer-Aided Designed Peptide Degrader Drug on PCSK9-Mediated Hypercholesterolemia

**DOI:** 10.1101/2025.06.25.661251

**Authors:** Gang Fan, Jingfen Lu, Ruirong Tan, Weiming Guo, Liang Hong, Jinhui Zha, Zhihan Tang, Jing Yang, Zhijian Yu, Miao Liu

**Author notes:** **Corresponding authors:** Miao Liu, M.D. & Ph.D., Department of Pathology, Brigham and Women’s Hospital Harvard Medical School Boston, MA 02115, United States.

## Abstract

Proprotein convertase subtilisin/kexin type 9 (PCSK9) is a key regulator of low-density lipoprotein receptor (LDLR) degradation, which leads to increased levels of low-density lipoprotein cholesterol (LDL-C) and a heightened risk of hypercholesterolemia. Existing therapeutic strategies—such as monoclonal antibodies and gene-editing approaches—are limited by high production costs, potential safety concerns, and limited effectiveness in targeting intracellular PCSK9. Targeted protein degradation (TPD) represents a promising alternative capable of overcoming these limitations. In this study, we utilized computer-aided drug design (CADD) to develop a peptide-based degrader, Cadd4, aimed at selectively reducing PCSK9 levels. Through molecular docking and structural refinement, a peptide sequence with strong binding affinity to PCSK9 was identified. Functional evaluation was conducted both in vitro using human LX-2 cells and in vivo in high-fat diet (HFD)-induced hypercholesterolemic mouse models. Cellular uptake, PCSK9 degradation, and LDLR restoration were assessed via confocal microscopy, western blotting, and biochemical analysis. Safety was evaluated by monitoring liver enzyme levels and general physiological parameters. Cadd4 exhibited efficient intracellular delivery and selective PCSK9 degradation in LX-2 cell. In HFD-fed mice, hepatic PCSK9 expression was reduced by 38%, accompanied by a marked increase in LDLR levels. These changes translated into a 25% reduction in total plasma cholesterol and a 29% decrease in LDL-C concentrations. Biodistribution studies indicated liver-specific accumulation of Cadd4, consistent with its intended site of action. Experiments using human liver tissue further confirmed its ability to decrease PCSK9 and enhance LDLR levels. No signs of liver toxicity or adverse systemic effects were observed throughout the study. This work introduces Cadd4 as a CADD-designed peptide-based degrader capable of lowering plasma cholesterol by promoting LDLR recovery via PCSK9 degradation. Its intracellular mechanism of action addresses key gaps in current PCSK9-targeted therapies and supports the broader utility of TPD platforms in managing hypercholesterolemia. These findings offer a foundation for developing cost-effective, scalable therapies within the context of precision cardiovascular medicine.

## Introduction

Proprotein convertase subtilisin/kexin type 9 (PCSK9) emerged as a critical therapeutic target for managing hypercholesterolemia in recent years [1]. By binding to low-density lipoprotein receptors (LDLR) and accelerating their degradation, PCSK9 significantly reduced the number of LDLR on the cell surface, thereby impairing the clearance of low-density lipoprotein cholesterol (LDL-C) by hepatocytes [2]. This mechanism resulted in elevated plasma LDL-C levels, which greatly increased the risk of atherosclerosis and cardiovascular diseases (CVD). Given its important role, developing effective therapies targeting PCSK9 became an essential focus for treating hypercholesterolemia [3, 4].

At the time, therapeutic strategies targeting PCSK9 included monoclonal antibodies (mAbs), tricyclic peptide inhibitors, and gene-editing technologies such as siRNA and CRISPR [5–7]. Among these, monoclonal antibodies were the most clinically advanced and widely used. Drugs such as evolocumab and alirocumab have demonstrated substantial efficacy in lowering low-density lipoprotein cholesterol (LDL-C) levels and reducing cardiovascular risk [7, 8]. However, these therapies are associated with several limitations. Their production and optimization processes are technically demanding and cost-intensive, restricting scalability and widespread access [9, 10]. Additionally, they function by neutralizing extracellular PCSK9 and preventing its interaction with the low-density lipoprotein receptor (LDLR), thereby inhibiting LDLR degradation [11]. This mechanism, however, does not address intracellular PCSK9 pools, potentially limiting their long-term efficacy. Moreover, delivery via injection lowers patient compliance and complicates chronic treatment regimens. Meanwhile, gene-editing approaches such as siRNA and CRISPR offer the theoretical advantage of long-term PCSK9 suppression at the genetic level. Despite this potential, these methods face significant barriers to clinical translation. Challenges include poor delivery efficiency, tissue toxicity (especially hepatic), and risks of off-target genetic modifications [12, 13]. In addition, the complexity of gene-editing systems and their associated development costs hinder their feasibility as broadly applicable treatments [13, 14]. These collective drawbacks highlight an urgent and unmet need for innovative, efficient, and cost-effective therapeutic modalities that can overcome the limitations of current PCSK9-targeted interventions.

Targeted protein degradation (TPD) technology has recently emerged as a highly promising therapeutic strategy for addressing PCSK9-related disorders. Unlike traditional functional inhibitors, TPD eliminates pathological proteins by actively inducing their degradation [15]. This mechanism not only overcomes the limitations of conventional therapies in terms of sustained inhibition and drug resistance but also opens new avenues for targeting previously "undruggable" proteins [16–19]. Among various TPD approaches, proteolysis-targeting chimeras (PROTACs) represent a distinct molecular class. By recruiting E3 ubiquitin ligases, PROTACs label the protein of interest (POI) for proteasomal degradation, enabling efficient and selective elimination of the target through a catalytic, event-driven process. This mechanism allows for continuous and thorough POI removal, offering the potential for a more durable therapeutic response. Most PROTACs developed to date are based on small molecules. However, designing small-molecule PROTACs for PCSK9 remains a major challenge. Due to the physicochemical properties of small molecules, it is often difficult to achieve high-affinity binding to large, structurally complex proteins like PCSK9, resulting in suboptimal degradation efficiency [20, 21]. Furthermore, the design of small-molecule PROTACs requires a delicate balance among biological activity, target selectivity, and pharmacokinetics. Combined with their complex synthetic routes and high development costs, these factors have significantly limited their applicability in targeting PCSK9.

Compared to small-molecule counterparts, peptide-based PROTACs exhibit a range of distinct and significant advantages. First, peptides generally possess low immunogenicity, which is of particular relevance in reducing the risk of immune-related adverse effects during clinical application [22, 23]. Second, conjugation with cell-penetrating peptides (CPPs) can substantially enhance membrane permeability, thereby enabling efficient intracellular delivery of the therapeutic payload to cytosolic targets [17, 18]. Furthermore, peptide synthesis technologies—particularly solid-phase peptide synthesis (SPPS)—are well-established and scalable, allowing for high-purity, customizable production suitable for industrial implementation at a reasonable cost [24, 25]. More importantly, peptides offer excellent structural tunability. Through rational sequence design, conformational control, and chemical modifications such as cyclization or PEGylation, peptides can be tailored to achieve high-affinity binding to specific targets such as PCSK9 [26]. These properties collectively endow peptide-based PROTACs with great potential for precise target engagement and controllable protein degradation. Nonetheless, these advantages come with certain technical challenges. The design and screening of bioactive peptides require robust technological infrastructure, particularly to ensure their functional potency, in vivo stability, and efficiency in inducing ubiquitin–proteasome system-mediated degradation. In recent years, the rapid development of computer-aided drug design (CADD) has provided critical support for the rational construction of peptide-based PROTACs. Utilizing techniques such as molecular docking, molecular dynamics simulations, and structural modeling algorithms, CADD enables rapid and accurate prediction of peptide–protein binding modes, significantly improving the efficiency and success rate of early-stage drug discovery and optimization efforts [27, 28]. Notably, peptide–target interactions often involve large, complex protein–protein interaction (PPI) interfaces, for which CADD exhibits superior predictive performance compared to its application in small-molecule ligand screening. As a result, CADD offers intrinsic advantages in the design of peptide-based PROTACs, making it a powerful tool for addressing traditionally "undruggable" targets.

In our previous work, we successfully developed a series of peptide-based PROTACs targeting diverse proteins with high degradation efficiency [27]. Building on this foundation, the present study employed CADD to develop a novel peptide-based degrader targeting PCSK9, aiming to address the limitations of existing therapeutic approaches. Through CADD, we achieved the precise design of peptide ligands with high binding affinity to PCSK9, and optimized their structural features via simulation-based screening to enhance both in vitro and in vivo degradation performance. Compared to conventional methods, CADD significantly improved screening efficiency and reduced development costs, making large-scale implementation more feasible. Functionally, multi-layered experimental validations demonstrated that the peptide degrader effectively reduced PCSK9 levels, restored LDLR expression, and enhanced LDL-C clearance, indicating strong therapeutic potential. This peptide-based targeted protein degradation strategy offers a novel and precise approach for the treatment of hypercholesterolemia and establishes a solid foundation for the future development of peptide-based PROTACs targeting other disease-relevant proteins.

PCSK9 plays a central role in the pathophysiology of hypercholesterolemia, and the limitations of current therapies—such as limited intracellular accessibility, and high manufacturing costs—underscore the need for innovative targeting strategies. By integrating CADD with peptide-based TPD technologies, this study presents an efficient, safe, and cost-effective therapeutic solution that not only expands treatment possibilities for hypercholesterolemia but also provides a theoretical and practical framework for the clinical translation and broader application of TPD platforms.

## Results

### 1. Design of Peptide Degrader and Membrane Penetration Ability

Using CADD, peptide ligands targeting PCSK9 were developed. Among the designed peptides, the top four candidates with the highest scores were synthesized into peptide degraders (Cadd1-4). Rhodamine-labeled peptide degraders were used to evaluate intracellular localization in LX-2 cells. Confocal laser microscopy demonstrated that with increasing concentrations and prolonged exposure, intracellular fluorescence intensity progressively increased, confirming the successful membrane penetration and intracellular localization of CADD-designed peptide degraders **(Figure 1 a-h)**. Western blot analysis showed that among the tested peptides, Cadd4 exhibited the most effective PCSK9 degradation **(Figure 1 i-p)**.

**Figure 1.**
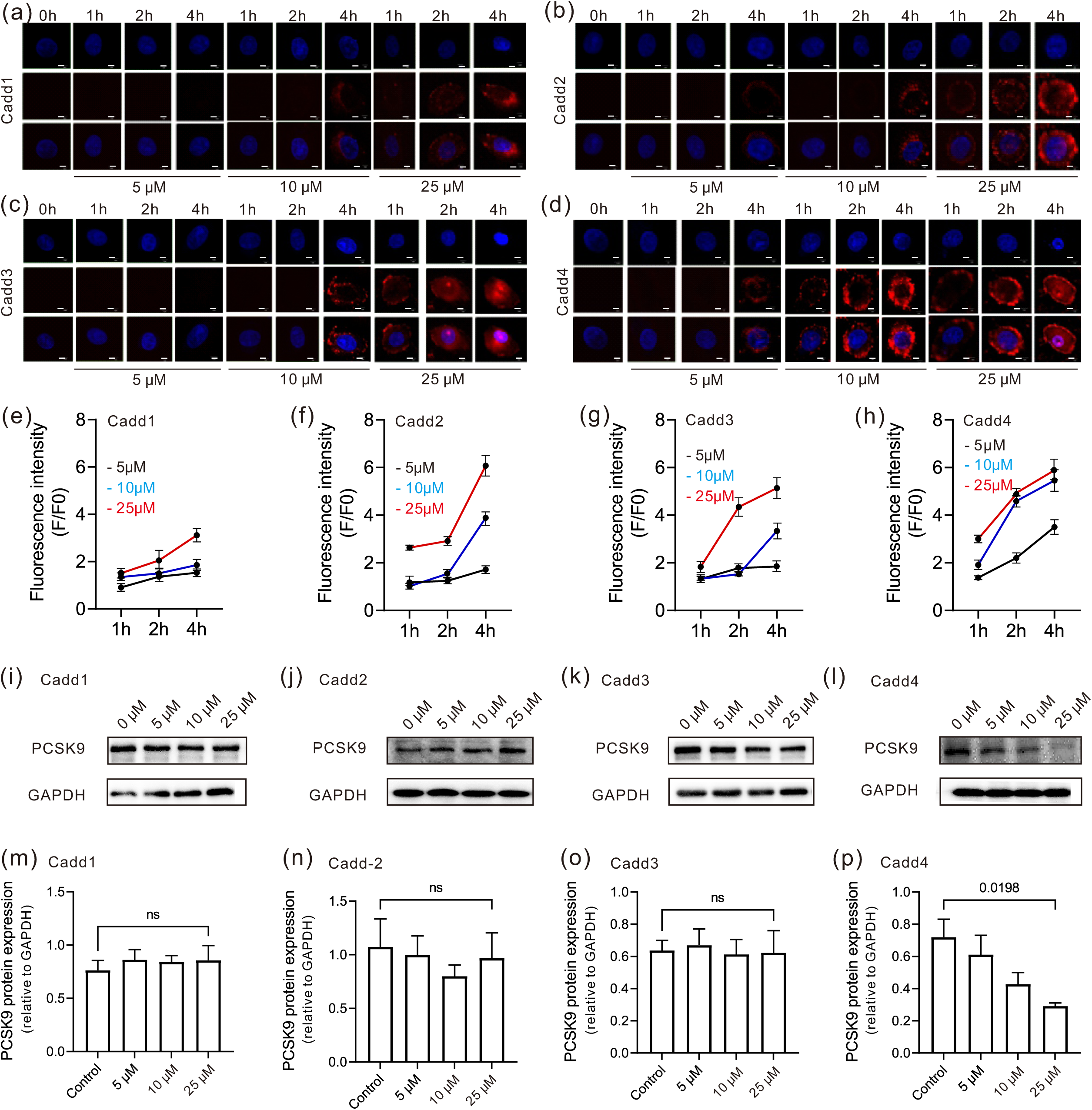
Computer-aided designed peptide degrader targeting of PCSK9. **(a-d)**, LX-2 cells were treated rhodamine-labeled Cadd 1-4 for 4 hours, the intracellular localization of Cadd 1-4 within LX-2 cells was examined through confocal laser microscopy. Red; Peptide-degrader Cadd4, blue; nucleus. (b-e, scale bar=5 μm). **(e-h)**, summary of the results (N=3). **(i-l)**, western blotting analysis of LX-2 cells after treatment with Cadd 1-4 the at the indicated doses and timepoints. **(m-p)**, quantification of western blot results (N=3). (Data were presented as means ± SD).

### 2. Cadd4-Induced PCSK9 Degradation

Cadd4 was selected for further in vitro and in vivo studies targeting PCSK9 **(Figure 2a)**. To elucidate its mechanism, Cellular Thermal Shift Assay (CETSA) was performed to evaluate Cadd4’s specific binding to PCSK9. The results revealed that Cadd4 significantly reduced the denaturation rate of PCSK9 at elevated temperatures, indicating a specific binding interaction **(Figure 2b)**. To determine whether the degradation of PCSK9 was mediated through the proteasome pathway, MG132, a proteasome inhibitor, was introduced. The results showed that MG132 effectively blocked the Cadd4-induced degradation of PCSK9, further confirming the proteasome-dependent degradation mechanism **(Figure 2c)**. Confocal microscopy demonstrated effective colocalization and binding between Cadd4 and intracellular PCSK9 **(Figure 2d)**.

**Figure 2.**
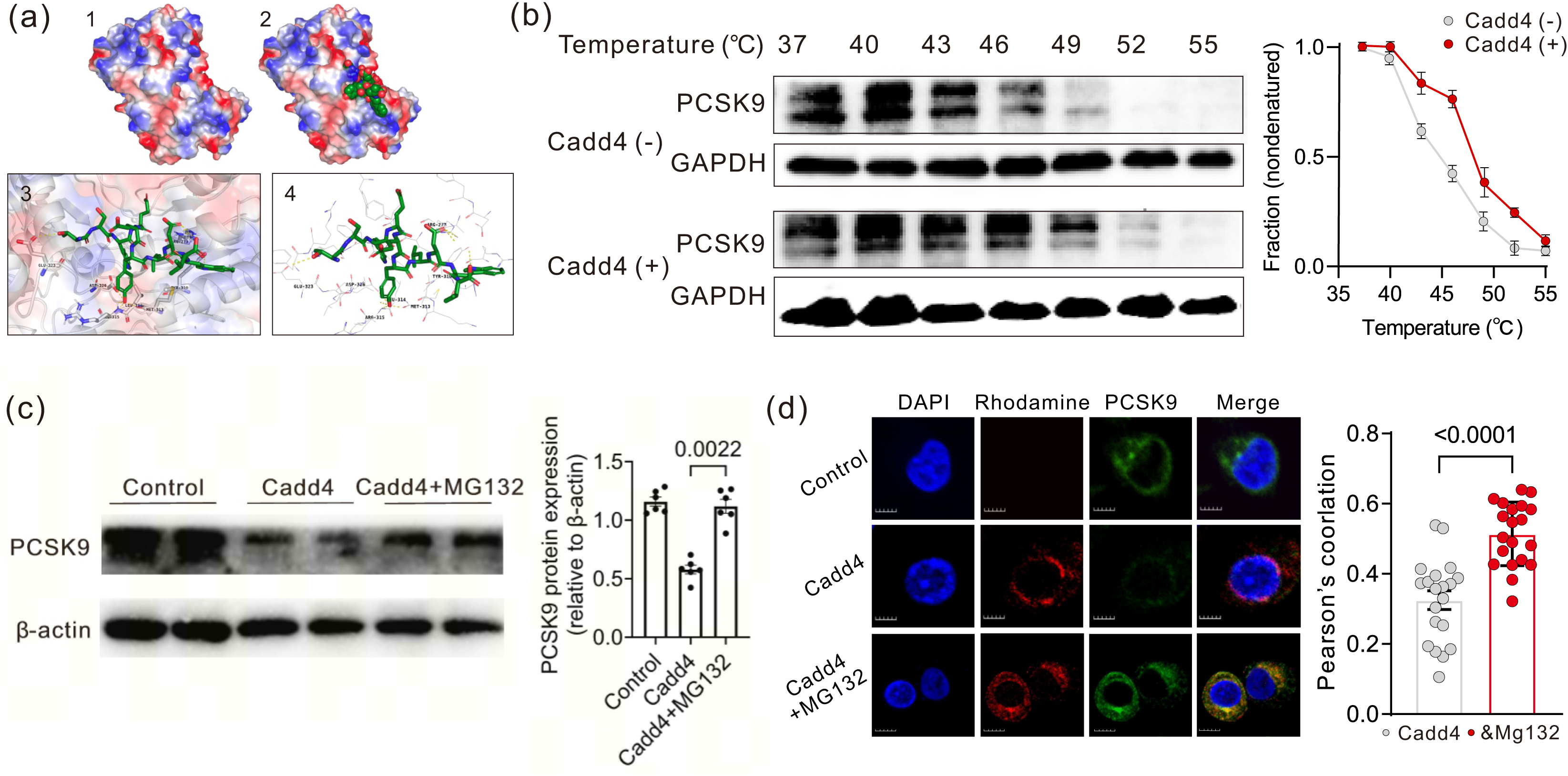
Cadd4 induces PCSK9 degradation through the proteasome pathway. **(a)**, Structural analysis of PCSK9-Cadd4 interaction: 1, Domain architecture of PCSK9; 2, Molecular docking of Cadd4 (green) into the catalytic site of PCSK9; 3-4, Crystal structure overlay generated using PyMOL. **(b)**, Cellular thermal shift assay (CETSA) demonstrating binding between Cadd4 and PCSK9. CETSA curves were derived from LX-2 cells treated with or without 20 μM Cadd4. GAPDH served as a loading control (N=3 replicates). **(c)**, western blotting analysis of 20 μM Cadd 4 treated LX-2 cells with and without MG132 (5 μM) (N=6). **(d)**, confocal microscopic analyses were employed to evaluate the impacts of 20 μM Cadd4 on PCSK9 expression, and its potential rescue by MG132 (5 μM) within LX-2 cells (N=3, n=5-10 cells/N) (Scale bars=10 μm) . Data were presented as means ± SD.

### 3. Distribution and Degradation Effects of Cadd4 In Vivo

The in vivo distribution of Cadd4 was evaluated using rhodamine-labeled Cadd4 in C57BL/6 mice. Both intraperitoneal (i.p.) and intravenous (i.v.) injections were tested. The results showed that Cadd4 rapidly localized to the liver with higher intensity and longer duration observed with i.p. injection **(Supplementary Figure 1)**. These findings suggest that i.p. administration was more effective for liver-specific targeting. Fluorescence imaging revealed that Cadd4 predominantly accumulated in the liver, reaching peak levels at 4 hours post-injection and declining significantly after 8 hours **(Figure 3 a-c)**. In high-fat diet (HFD)-induced hypercholesterolemic mice, Cadd4 reduced hepatic PCSK9 expression by approximately 38% and concurrently upregulated LDLR expression **(Figure 4 a-b)**. Tissue section analysis further confirmed that Cadd4-induced PCSK9 degradation led to a marked increase in hepatic LDLR levels **(Figure 4 c)**.

**Figure 3.**
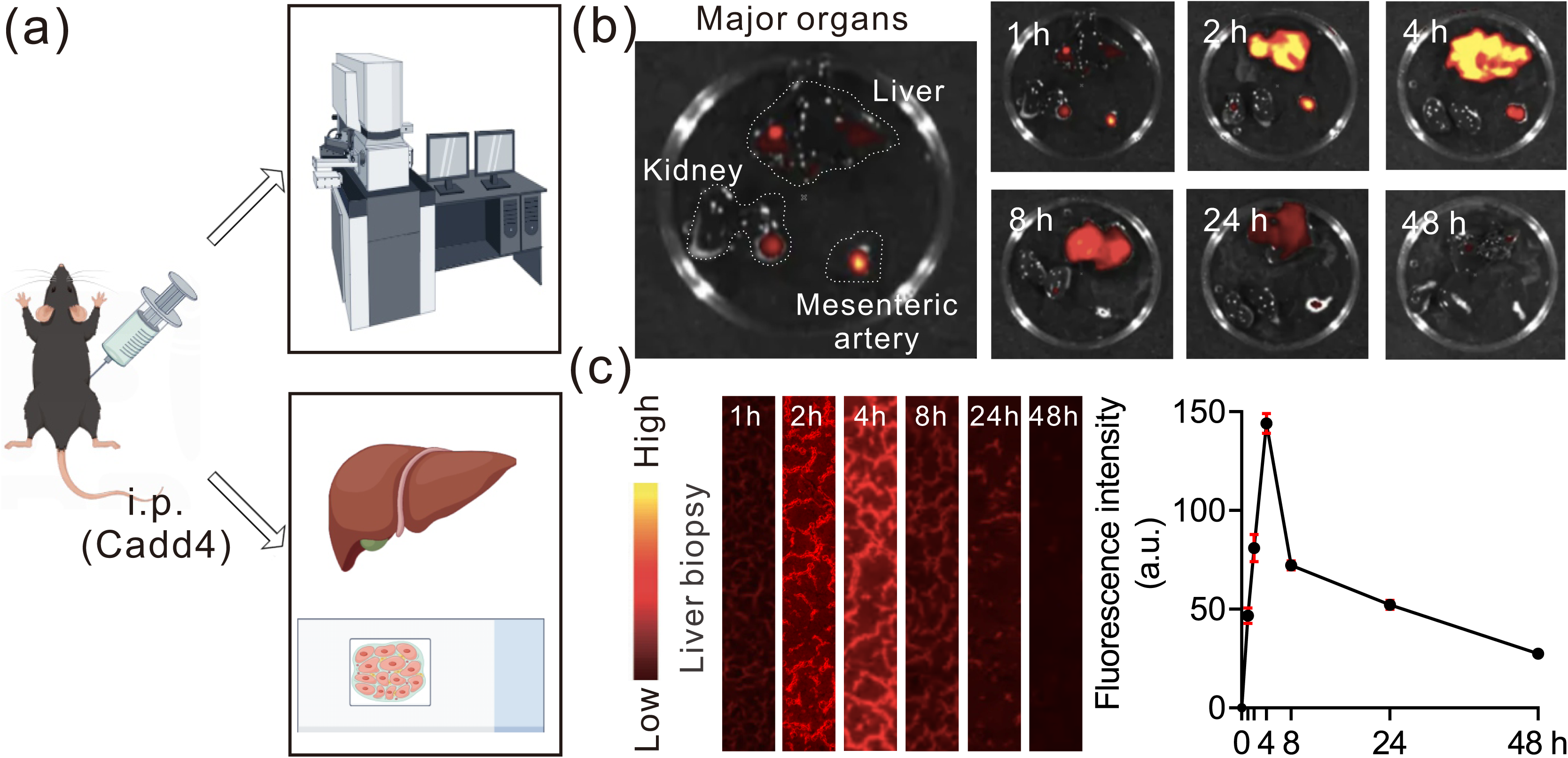
Biodistribution and circulation lifetime of Cadd4 in vivo. **(a-c)**, fluorescence imaging of the biodistribution and liver biopsy of Cadd4 in C57BL/6 mice at a series of time points, and quantitative biodistribution analysis of rhodamine-labeled Cadd4 in liver biopsy (N=3). Data were presented as means ± SD.

**Figure 4.**
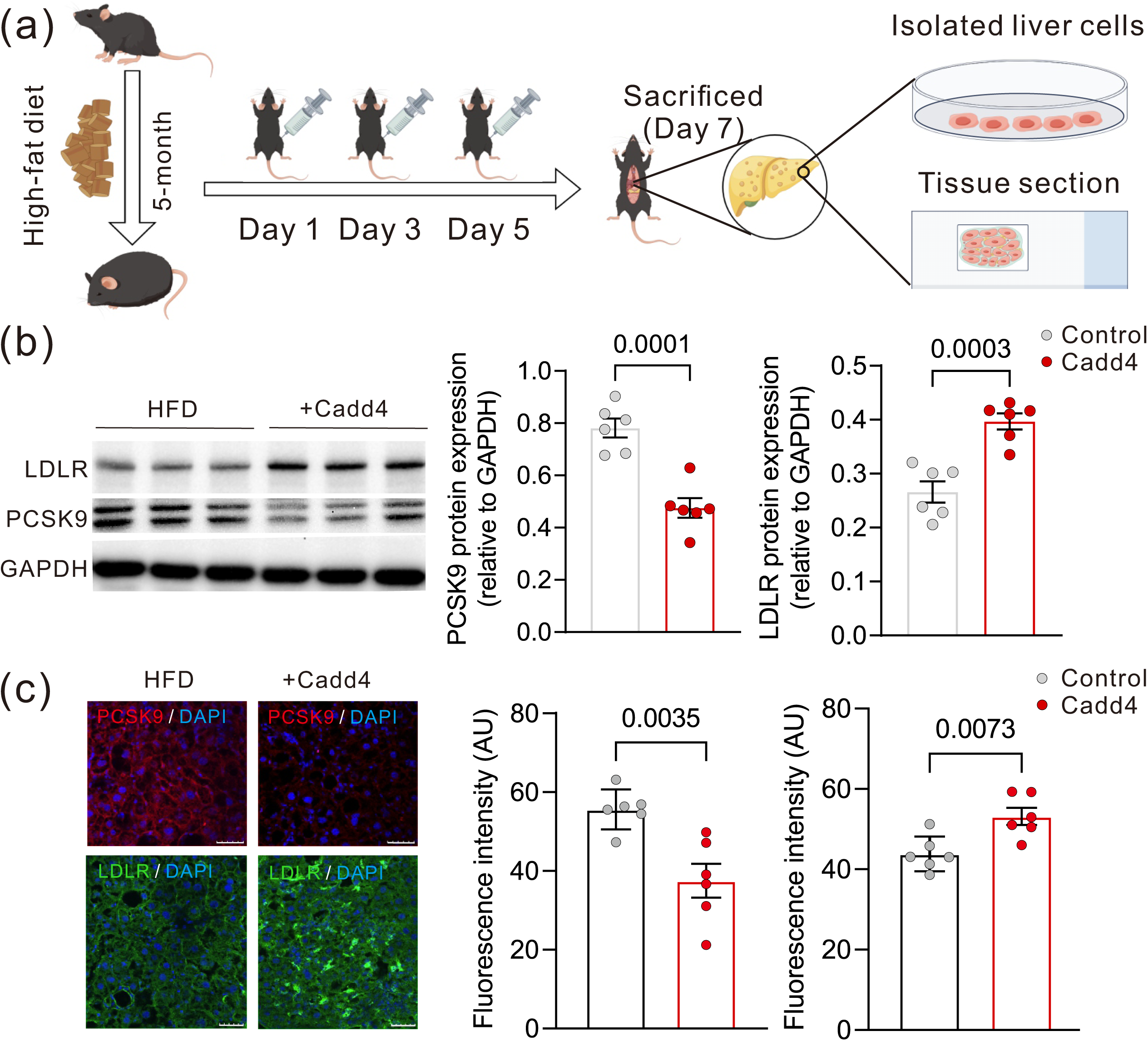
Cadd4 induces PCSK9 degradation in vivo. **(a)**, experimental design of Cadd4 administration in HFD mice is shown. **(b)**, western blotting analysis of PCSK9 protein and LDLR protein in HFD mice treated with and without PCSK9 degrader Cadd4 (25 μM/kg) (N=6). **(c)**, confocal microscopic analyses were employed to evaluate the level of PCSK9 protein and LDLR protein in mice (N=6)(Scale bars=100 μm). (Data were presented as means ± SD).

### 4. Effects of Cadd4 on Lipid Metabolism

The effects of Cadd4 on lipid metabolism were assessed in HFD-fed mice and ApoE-deficient mice **(Figure 5 a, Supplementary Figure 2 a)**. Compared to the control normal diet (ND) mice, the HFD group exhibited a significant elevation in plasma total cholesterol (TC) levels, from 2.0 mM to 5.5 mM, and in LDL-C levels, from 0.40 mM to 1.7 mM. Remarkably, administering Peptide-degrader three times over a 7-day period significantly reversed the metabolic changes in mice fed a high-fat diet, showing a 25% reduction in TC and a 29% decrease in LDL-C **(Figure 5 b).** Additionally, studies in ApoE-deficient mice supported the cholesterol-lowering effects of Cadd4, demonstrating significant decreases in TC and LDL-C levels (**Supplementary Figure 2b)**.

**Figure 5.**
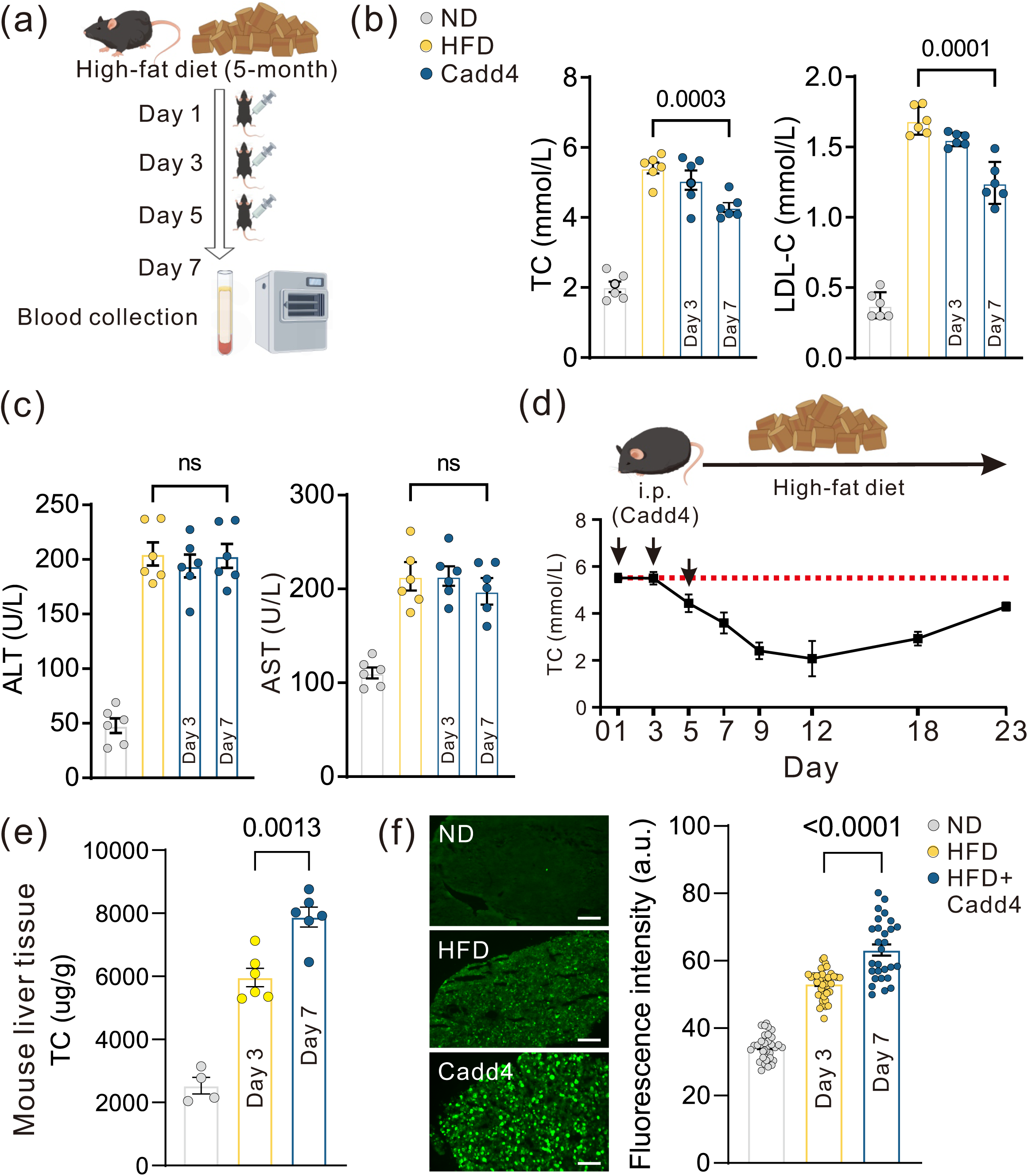
Therapeutic peptide-degrader against plasma cholesterol. **(a)**, experimental design of Cadd4 administration in HFD mice is shown. **(b-c)**, serum levels of TC, LDL-C, TP and AST in the ND mice, HFD mice treated with and without PCSK9 degrader Cadd4 (N=6). **(d)**, serum TC analyses were employed to evaluate the duration of Cadd4 in vivo (N=5). **(e)**, quantitative liver TC value by using Elisa kit (N=6). **(f)**, lipid droplets green fluorescence assay, and quantitative intensity analysis of lipid droplets green fluorescence in liver biopsy (N=6, n=3-8 sections/N)(Scale bars=500 μm). (Data were presented as means ± SD).

Additionally, we also quantified the liver steatosis in HFD mice with and without Cadd4 treatment. Our findings indicate a significantly increase in TC value in HFD mice treated with Peptide-degrader Cadd4 (**Figure 5 e**). These observations aligned with the results obtained from lipid droplets green fluorescence assay, which demonstrated an increased presence of lipid droplets upon Cadd4 treatment (**Figure 5 f**). These data suggest that the degradation of PCSK9 proteins, prompted by Cadd4, is linked to the increasing of LDLR levels, thus contributing to the increased TC accumulation in the liver.

### 5. Effects of Cadd4 in Human Liver Tissue

To evaluate the efficacy of Cadd4 in human tissues, experiments were performed on liver samples obtained from surgical resections **(Figure 6 a)**. Results showed that rhodamine-labeled Cadd4 successfully penetrated cell membranes and localized within liver cells. Western blot and immunostaining analyses further demonstrated effective PCSK9 degradation and significant upregulation of LDLR expression **(Figure 6 b-c)**.

**Figure 6.**
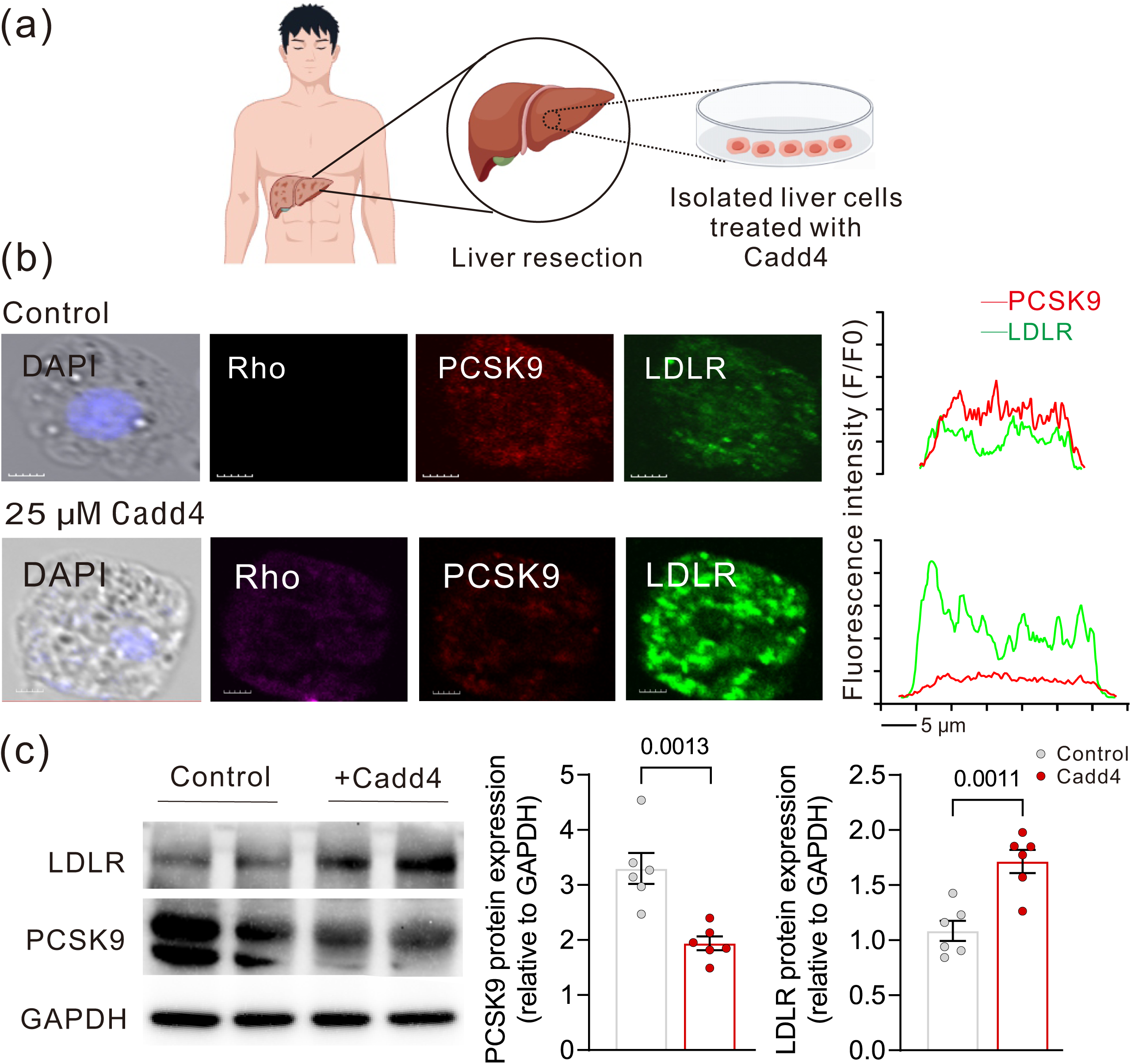
Cadd4 induces PCSK9 degradation in human liver. **(a)**, experimental design of Cadd4 administration in human liver is shown. **(b)**, in situ hybridization of rhodamine-labeled Cadd4 (purple), PCSK9 (red) and LDLR (green) in the isolated human liver cells. Nuclei are visualized with DAPI. (Scale bar=5 μm). **(c)**, western blotting analysis of PCSK9 protein and LDLR protein in isolated human liver cells treated with and without PCSK9 degrader Cadd4 (N=6). (Data were presented as means ± SD).

### 6. Safety Evaluation

Safety evaluation revealed that Cadd4 treatment had no significant effects on mouse body weight, food intake, or liver function indicators (ALT and AST levels), indicating excellent safety profiles **(Figure 5 c)**.

## DISCUSSION

This study developed a peptide-based PCSK9 degrader, Cadd4, using CADD technology, and systematically validated its strong degradation activity, and favorable safety profile through both in vitro and in vivo experiments. Multiple PCSK9-targeting therapeutic strategies have been proposed, including antibody drugs such as Evolocumab and Alirocumab, as well as gene-editing approaches like siRNA and CRISPR [5–7, 29]. Although these treatments have shown some effectiveness in lowering LDL-C levels, they still face practical challenges. For example, antibodies mainly target circulating PCSK9, with limited ability to affect intracellular protein, which restricts their applicability under certain pathological conditions [30]. Gene-editing methods, while offering theoretical potential for long-term suppression, remain limited in clinical translation due to safety concerns, off-target effects, and delivery inefficiencies [12, 13, 31].

Cadd4’s molecular mechanism of PCSK9 degradation was validated through integrated cellular assays, confirming its proteasome-dependent action via MG132 inhibition and direct target engagement via CETSA, which stabilized PCSK9-Cadd4 interactions under thermal stress. Confocal microscopy demonstrated intracellular colocalization of Cadd4 with PCSK9, while Western blotting revealed dose-dependent PCSK9 reduction (50% at 20 µM, ∼90% at 50 µM) and concomitant LDLR upregulation, mechanistically linking degradation to functional LDL-C clearance. The absence of PCSK9 recovery post-MG132 treatment underscores the UPS-driven, irreversible nature of this process, distinguishing it from transient inhibitory modalities [27, 32]. These findings collectively elucidate Cadd4’s dual role as a PCSK9 degrader and LDLR restorer, bridging molecular targeting to therapeutic efficacy in lipid homeostasis.

Cadd4’s therapeutic superiority over existing PCSK9-targeting modalities stems from its PROTAC-mediated degradation mechanism, which addresses key limitations of conventional approaches. Unlike traditional PCSK9 inhibitors - including small-molecule antagonists and mAbs that temporarily inhibit enzymatic activity through competitive binding [11, 33, 34], our peptide-based degrader induces sustained PCSK9 depletion via ubiquitin-proteasome system engagement, as evidenced by eliminating 38% of hepatic PCSK9 protein and lowering plasma LDL-C by 29% in HFD mice. This complete PCSK9 removal-contrasting with partial inhibition by small-molecule antagonists and mAbs-ensures durable LDLR restoration and minimizes compensatory PCSK9 accumulation risks. Mechanistically, while mAbs merely block extracellular PCSK9-LDLR interactions, Cadd4’s ubiquitin-proteasome-driven degradation eradicates both intracellular and secreted PCSK9 pools, offering broader target coverage. Structurally, Cadd4’s CADD-optimized linear peptide design simplifies synthesis compared to cyclization-dependent inhibitors, enhancing scalability and cost-effectiveness-a critical advantage over mAbs, which require expensive mammalian cell production ($5,000–14,000 annually). Additionally, peptide PROTACs exhibit lower immunogenicity, circumventing antibody resistance risks, though their proteome-wide degradation impacts warrant further study [22, 23]. By integrating computational design with catalytic efficiency, Cadd4 bypasses the genomic instability concerns of gene-editing technologies while outperforming single-target therapies in lipid-lowering durability. These attributes position it as a transformative, cost-accessible alternative in hypercholesterolemia management, bridging gaps left by current biologics and small-molecule inhibitors. Furthermore, Cadd4-induced PCSK9 degradation directly enhances LDLR recycling by blocking PCSK9-mediated lysosomal targeting, resulting in sustained hepatic LDLR upregulation and a reduction in plasma LDL-C, as observed in HFD mice. While prolonged LDLR elevation could theoretically activate feedback mechanisms (e.g., SREBP2 suppression due to hepatic cholesterol accumulation), no significant LDLR downregulation was detected during treatment, suggesting Cadd4’s dominant short-term efficacy over compensatory pathways [35, 36]. Unlike broad-spectrum lipid modulators, Cadd4’s specific PCSK9 targeting minimizes off-target effects on triglyceride-rich lipoprotein metabolism, as PCSK9 primarily regulates LDLR rather than LRP1 or VLDLR [37, 38]. Additionally, by avoiding direct HMGCR inhibition, Cadd4 sidesteps the compensatory cholesterol synthesis triggered by statins, potentially reducing adverse effects linked to dysregulated biosynthesis. These findings highlight Cadd4’s precision in reshaping lipid homeostasis while maintaining a favorable risk-benefit profile for LDL-C management.

Previous studies have systematically characterized diverse classes of peptide-based PCSK9 inhibitors, including nature-derived peptides, cyclic peptides from combinatorial libraries, and peptidomimetics [39–42]. These agents primarily function by competitively blocking the PCSK9–LDLR interaction and have demonstrated promising efficacy in preclinical models. In contrast, Cadd4 induces the proteasome-dependent clearance of PCSK9, which enables comprehensive depletion of both intracellular and extracellular PCSK9 pools. This mechanism allows for more durable restoration of LDLR activity and offers therapeutic advantages over conventional blockade-based strategies. By leveraging a degradative mode of action, Cadd4 expands the conceptual landscape of peptide-based interventions for hypercholesterolemia.

CADD played a key role in the design and development of Cadd4 throughout this study. By employing molecular docking and structural optimization, CADD facilitated the screening of multiple potential PCSK9 recognition sequences and refined the binding mode of the degrader with PCSK9’s active sites. The final selection, Cadd4, exhibited excellent binding affinity and structural stability, providing a high-quality candidate for subsequent experimental validation. Cadd4’s performance in HFD and ApoE^-/-^ models further supports its broad applicability in cholesterol reduction. Experimental results showed that Cadd4 significantly reduced plasma cholesterol levels and upregulated LDLR expression in both models, confirming its suitability. In HFD-induced hypercholesterolemic mice, Cadd4’s liver-specific distribution and cholesterol-lowering effects were particularly prominent. Gene-editing therapies, such as CRISPR, rely on delivery systems to achieve liver specificity, while Cadd4 achieves hepatic enrichment through simple intraperitoneal injection, avoiding the complexity of delivery systems and reducing off-target effects. Additionally, unlike other therapies with prolonged systemic metabolism, Cadd4’s rapid clearance profile enhances safety. Fluorescence imaging demonstrated that Cadd4 reached peak liver levels within 4 hours post-injection and significantly decreased by 8 hours, minimizing drug accumulation in non-target organs and reducing potential toxicity. Importantly, the absence of a hook effect in Cadd4, despite its prevalence in conventional degraders [43], may arise from its computer-optimized design and peptide-based structural flexibility, which stabilize ternary complex formation and minimize non-productive binary interactions at high concentrations. Controlled cellular uptake dynamics, distinct from small molecules, likely prevent intracellular saturation, sustaining degradation efficiency across dose ranges. These attributes, corroborated by stable in vitro and in vivo activity, highlight Cadd4’s enhanced pharmacokinetic controllability and translational promise. Existing therapies have shown cross-species applicability in preclinical studies but lack direct validation in human liver tissue [7–9, 44]. This study demonstrated, for the first time, that Cadd4 effectively degrades PCSK9 and restores LDLR expression in human liver samples, with ongoing preclinical optimization to enhance pharmacokinetic stability and safety profiles for clinical translation. Future efforts will focus on structural modifications (e.g., PEGylation, cyclization) to improve oral bioavailability, extended toxicological assessments in non-human primates, and evaluation of synergistic potential with statins to address the limitations of current injectable PCSK9 inhibitors, such as high costs and poor adherence.

Although Cadd4 exhibits promising therapeutic potential, several challenges remain in its clinical translation. One key issue lies in the susceptibility of its peptide structure to enzymatic degradation in vivo, which may necessitate chemical modifications such as PEGylation or cyclization to enhance metabolic stability and improve oral bioavailability. In addition, future studies will focus on comprehensive toxicological evaluations in non-human primates and explore its potential for combination therapy with statins to further expand its clinical utility. The HFD-induced C57BL/6 mouse model is commonly employed in the study of lipid metabolism; however, its applicability to human pathology warrants critical examination. Although this model replicates certain hallmarks of hypercholesterolemia, including elevated plasma LDL-C and total cholesterol, interspecies differences exist. Notably, mice predominantly transport cholesterol via HDL, whereas humans rely more heavily on LDL [45, 46]. Nevertheless, mice subjected to high-fat feeding exhibit increased LDL-C levels due to suppression of hepatic LDLR expression, a mechanism reminiscent of PCSK9-mediated LDL-C accumulation in humans. The regulatory interaction between PCSK9 and LDLR appears to be well conserved in this model . In our study, administration of the test compound led to a 38% reduction in hepatic PCSK9 levels, accompanied by an upregulation of LDLR, consistent with the pharmacodynamic effects observed in clinical settings involving PCSK9-targeting therapeutics. Despite these advantages, atherosclerotic lesions in high-fat diet-fed mice develop more slowly and with reduced severity compared to humans, limiting its predictive value for long-term cardiovascular outcomes. To address this concern, we extended our investigation to human liver tissue, where we observed comparable reductions in PCSK9 and restoration of LDLR expression following treatment. This cross-species consistency supports the relevance of our findings to human physiology.

To comprehensively evaluate the safety, specificity, and selectivity of Cadd4, we employed a range of validation methods aligned with our prior TPD research, including cell membrane permeability assessment, and in vivo toxicological analysis [27, 32]. Importantly, peptide-based PROTACs function exclusively at the protein level without interacting with genomic DNA, thereby minimizing genotoxic risk compared to gene-editing technologies [27, 32, 47, 48]. In vivo safety indicators—including serum ALT and AST levels, body weight, and food intake—remained within normal ranges throughout the study. These results collectively support the favorable safety, metabolic stability, and target selectivity of Cadd4. Nevertheless, further investigation is needed to define the dose-response relationship and assess long-term toxicity through large-scale animal studies and eventual clinical trials.

## CONCLUSIONS

In this study, a novel peptide degrader, Cadd4, was successfully designed and constructed using CADD technology. Its potent ability to target and degrade PCSK9 was validated, along with its favorable liver-targeting characteristics. By accelerating the degradation of PCSK9, Cadd4 effectively upregulated LDLR expression and improved cholesterol metabolism. A series of in vitro and in vivo experiments confirmed Cadd4’s excellent performance in cellular uptake, target specificity, and safety across multiple dimensions. This work not only broadens the application scope of CADD in drug development but also provides a theoretical foundation for innovative therapeutic strategies against hypercholesterolemia. Future efforts will focus on enhancing the stability of Cadd4 and optimizing its manufacturing process to facilitate clinical translation.

## MATERIALS AND METHODS

### 1. Computational Design and Synthesis of a PCSK9-Targeted Peptide Degrader

To develop a peptide-based degrader targeting PCSK9, we adopted a structure-guided computational strategy in combination with chemical synthesis and experimental validation. This workflow builds upon our previously established CADD platform [27]. Based on this framework, we designed a series of 10-mer peptide sequences aimed at high-affinity binding to PCSK9. The crystal structure of PCSK9 was obtained from the Protein Data Bank (PDB ID: 4NMX), as previously reported by Zhang et al.[49]. Peptide–protein docking was performed using the HPEPDOCK server. Docking simulations were conducted using default parameters, generating multiple conformations for each peptide. These conformers were initially screened based on their predicted binding affinity, spatial complementarity, and ability to interact with key residues within the PCSK9 binding interface. To enhance the rigor of the selection, we employed the MODPEP tool to generate diverse conformational ensembles of each peptide prior to docking. The MDock scoring function was used for evaluation, incorporating van der Waals forces, electrostatic interactions, and geometric alignment. Final sequences were selected based on a combination of the lowest predicted binding energy and the highest structural stability. In addition to global scoring metrics, we paid particular attention to residue-level interaction patterns, specifically the formation of hydrogen bonds and hydrophobic contacts that contribute to stable peptide–protein binding.We selected the peptide with the highest predicted binding affinity—TSWEEYLDWV—for synthesis, based on computational docking results. This sequence demonstrated superior binding among all candidates and was chosen as the lead for further development. Its PCSK9 degradation efficacy was subsequently confirmed in cell-based assays. To generate a functional peptide-based degrader, we conjugated this sequence to two essential components: a rhodamine fluorophore for fluorescence-based visualization, and the ALAPYIP motif, a well-characterized ligand for recruiting the Von Hippel-Lindau (VHL) E3 ubiquitin ligase complex. The linker bridging the PCSK9-binding peptide and the E3 ligase ligand plays a critical role in maintaining both structural flexibility and functional independence. In this study, we utilized the GSGS linker sequence, which is widely recognized in peptide PROTAC designs for its stability and efficiency. The selection of both the linker and E3 ligase ligand was grounded in established literature and empirical data. As peptide-based PROTACs typically involve constrained linker options to preserve conformational freedom and bioactivity, the GSGS linker was chosen for its consistent performance in prior studies. Likewise, the ALAPYIP-based VHL ligand was selected due to its proven reliability across multiple reports, which have demonstrated its ability to achieve robust and selective degradation in peptide PROTAC systems [50–52]. Taken together, these design choices reflect a rational, evidence-based strategy to ensure optimal degradation efficiency and structural integrity in the engineered degrader. All final peptides were synthesized by SynPeptide Co. Ltd (Nanjing, China). with a purity exceeding 95%, confirmed by high-performance liquid chromatography (HPLC). The identity of each compound was verified by electrospray ionization mass spectrometry (ESI-MS), as presented in Supplementary Data 1 and 2. This work establishes a rational and efficient pipeline for engineering peptide-based degraders capable of targeting proteins such as PCSK9.

### 2. Animals

In total, 8- to 12-week-old male C57BL/6 mice were used. All animals were maintained at the breeding facility of the Animal Centre of Shenzhen University Affiliated Sixth Hospital in individually ventilated cages under standardized conditions that included a 12-hour dark-light cycle and free access to drinking water. Mice were fed a regular chow diet (Normal diet, ND) or a high-fat diet (HFD) (Catalogue, ASHF4, Dyets) during the study. B6J-Apoe-KO mice (Strain NO. C001507) were purchased from Cyagen. All animals were deeply anaesthetized by inhalation of isoflurane until cessation of breathing, then killed by cervical dislocation and the organs removed. All animal protocols were approved by the local animal care committee of Shenzhen University Affiliated Sixth Hospital and the ethics committee of TOPBIOTECH Shenzhen (reference number: TOP-IACUC-2023-0256). There are no ethical concerns.

### 3. Human samples and Ethical approval

Human liver samples were extracted from resected liver tissue. The trial was conducted in accordance with the Declaration of Helsinki and the International Conference on Harmonisation Good Clinical Practice Guidelines and the protocol was approved by the ethics committee of Shenzhen University Affiliated Sixth Hospital (reference number: CTR20221529).

### 4. Cell lines and cell culture

Human hepatic stellate cell line, LX-2, was been described previously [53]. LX-2 cells were cultured at 37 °C in a humidified atmosphere containing 5% CO2 with Dulbecco’s modified Eagle’s medium (DMEM) containing 10% fetal bovine serum, 100 U of penicillin and 100 μg of streptomycin (all from Gibco BRL, CA).

### 5. Western blot analysis

Following treatment, protein lysates were obtained from cultured cells or isolated cells, and western blotting was performed according to a standard protocol [27]. Briefly, cells were collected in EP tubes, lysed in RIPA buffer, and the protein concentration of cell lysates were measured by Bradford assay. Equal quantities of protein were separated by 10% SDS-PAGE and then transferred onto a nitrocellulose membrane. After blocking the membrane in 5% skim milk at room temperature, the membrane was incubated with primary antibody as follows: anti-PCSK9 rabbit polyclonal antibody (#55206-1-AP, 1:500, Proteintech, China); anti-LDLR rabbit mmonoclonal aantibody (#AG2503, 1:500, Beyotime, China) and internal control GAPDH, prepared in 1% BSA in TBST at 4 °C overnight. Next, the membrane was washed and stained with secondary antibody for 2 h at room temperature. Signals were then detected using an ECL system (Vazyme, Jiangsu, China) and a ChemiDoc XRS imaging system (Bio-Rad Laboratories Inc., Heracles, CA, USA), and images were analyzed by ImageJ.

### 6. Cellular thermal shift assay (CETSA)

Cells were cultured and treated with the Cadd4 at 20 µM for 4 hours, then were collected into PCR-tubes, pelleted by centrifugation, and, after careful removal of the supernatant, resuspended in PBS [27]. The tubes were kept at room temperature until the heat treatment step, which involved heating the PCR-tube strips to their designated temperature (seven temperature endpoints between 37 and 55 °C) for 3 min (Veriti thermal cycler, Applied Biosystems), then cooling for 3 min at room temperature. The cell pellet was lysed with RIPA buffer at 4 °C for 30 min and centrifuged at 20000 X g for 20 min at 4 °C. The detection and quantification of the soluble protein was then achieved by western blotting.

### 7. Biodistribution of Cadd4 by In Vivo Imaging System

Biodistribution analysis using vivo imaging system was been described previously [54]. The rhodamine-labeled Cadd4 were intraperitoneal injected into the C57BL/6 mice. The mice were anesthetized with 2% isoflurane, and their distribution was monitored at pointed times (1, 2, 4, 8, 24, and 48 h, respectively) by an imaging system (InSyTe FLECT/CT). After euthanasia of the mice, the dissected major organs (liver, kidney and mesenteric artery) were collected for ex vivo imaging. The fluorescence intensity of the isolated organs was quantified using ImageJ.

### 8. Histological analyses

Mice were anesthetized with 2% ketamine/10% rompun, perfused by 30 ml PBS and 50 ml 4% PFA (Roth, diluted in PBS) and after wards liver were dissected, and tissue pieces were further fixed for 4 h in 4% PFA, transferred to 15% sucrose (in PBS, Merck) for 4 h and incubated in 30% sucrose overnight. After embedding in TissueTek (Sakura), the tissue is frozen at -80 °C and 8 µm sections were obtained in a Leica cryostat at -30 °C. For immunostainings, the tissue pieces/cells were incubated with blocking buffer (1% donkey serum/1% TritonX100/PBS), the first antibody was applied overnight at 4°C, and after washing with PBS/1%Tween, the secondary antibody and DAPI were applied for 2 h. Afterwards the tissue pieces/cells were embedded in ImmoMount (ThermoScientific #9990402). Antibodies: anti-PCSK9 rabbit polyclonal antibody (55206-1-AP, 1:100, Proteintech, China), anti-LDLR rabbit mmonoclonal aantibody (AG2503, 1:100, Beyotime, China), DAPI (#D9542, Sigma). Boron Dipyrromethene (BODIPY, Invitrogen) was used to detect lipid accumulation. The stained sections were analysed with confocal microscope (FV3000, Olympus), and images were analysed by ImageJ.

### 9. Biochemical assays

Whole blood samples were collected via retro-orbital plexus from each mouse. Following collection, the blood was allowed to undergo the clotting process at room temperature for a duration of 30 minutes. Subsequently, the resulting serum was isolated through centrifugation at a force of 1,500g for a time period of 10 minutes. The concentrations of TC and serum LDL-C within the serum were quantified using the TC assay kit as well as the LDL-C assay kit sourced from Beyotime in China. Lipids present in the liver were initially extracted through a methanol-chloroform composition (at a ratio of 2:1, v/v). The hepatic TC contents were then determined utilizing commercially available assay kits from Shanghai Kehua bio-engineering Co. The levels of circulating ALT and AST were evaluated employing commercial kits procured from Beyotime in China.

### 10. Statistical analysis

All experimental data were presented as means ± standard deviation (means ± SD). The Shapiro-Wilk normality test was applied to assess the normal distribution of the data. For normally distributed data, statistical comparisons between two groups were performed using an unpaired Student’s t-test, while one-way ANOVA followed by Bonferroni’s multiple comparison test was used for comparisons involving three or more groups. For datasets with non-normal distributions, nonparametric tests were employed: the Mann-Whitney U test for two-group comparisons and the Kruskal-Wallis test with Dunn’s post hoc analysis for multiple groups. Pearson’s correlation coefficient for normally distributed data. Sample sizes are explicitly indicated in figure legends. All statistical analyses were performed using GraphPad Prism (version 9.0) .

## Author contributions

All authors were responsible for interpretation of the data, contributed to the drafting and revised the manuscript critically for important intellectual content. G.F., Z.Y., and M. L. designed the project and M.L. developed the peptides. All authors have approved the final version of the manuscript and agree to be accountable for all aspects of the work. All persons designated as authors qualify for authorship, and all those who qualify for authorship are listed.

## Funding

This work was supported by the National Natural Science Foundation of China (82001489), Guangdong Natural Science Foundation (2020A1515110158), Shenzhen Medical Research Fund (A2406066), Shenzhen Natural Science Foundation (JCYJ20220530141613031), Shenzhen Nanshan District Science and Technology Plan Project (NS044) and Shenzhen University Affiliated Sixth Hospital Medical Research Foundation.

### Availability of data and materials

The datasets used and/or analysed during the current study are available from the corresponding author upon reasonable request.

## Declarations

### Ethics approval and consent to participate

The study was reviewed and approved by the ethics committee of Shenzhen university Affiliated Sixth Hospital (reference number: CTR20221529) and the ethics committee of TOPBIOTECH Shenzhen (reference number: TOP-IACUC-2023-0256). There are no ethical concerns.

### Consent for publication

Not applicable.

---

**Supplementary figure 1.**
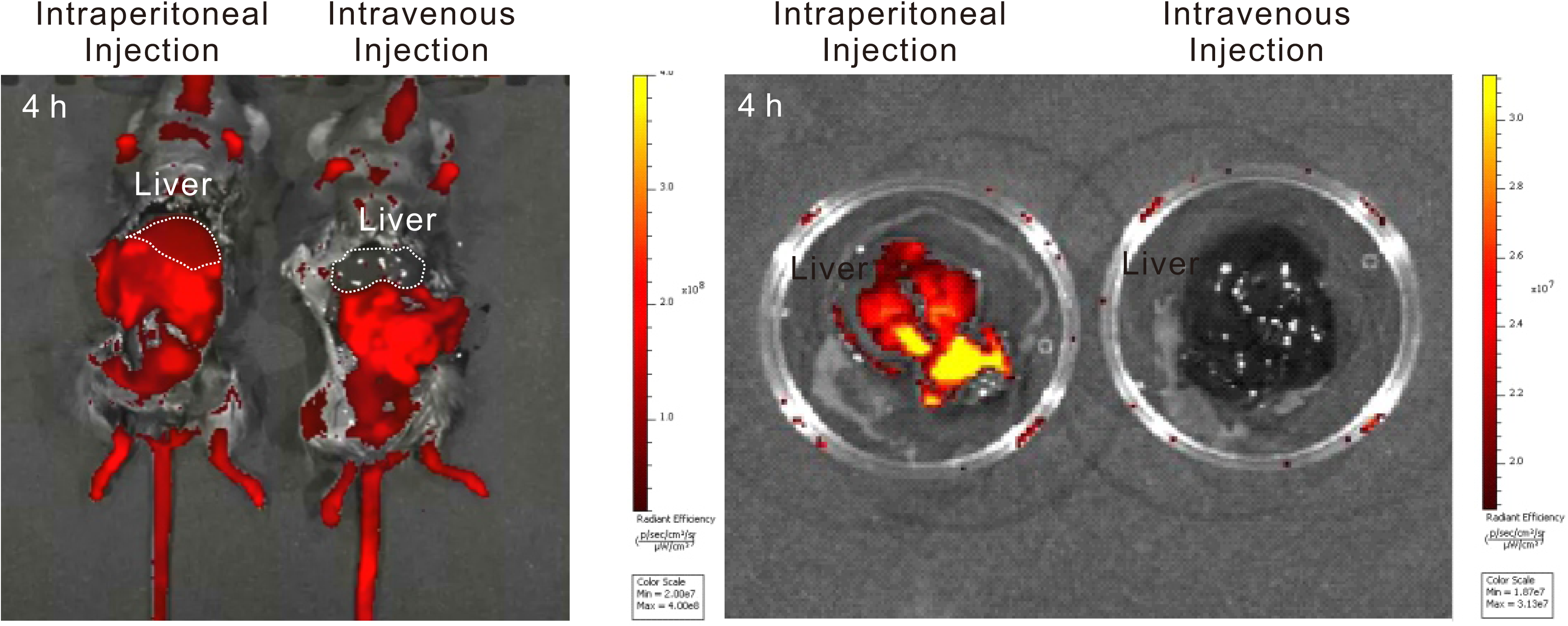
Intraperitoneal injection and intravenous injection of Cadd4 in mice.

**Supplementary figure 2.**
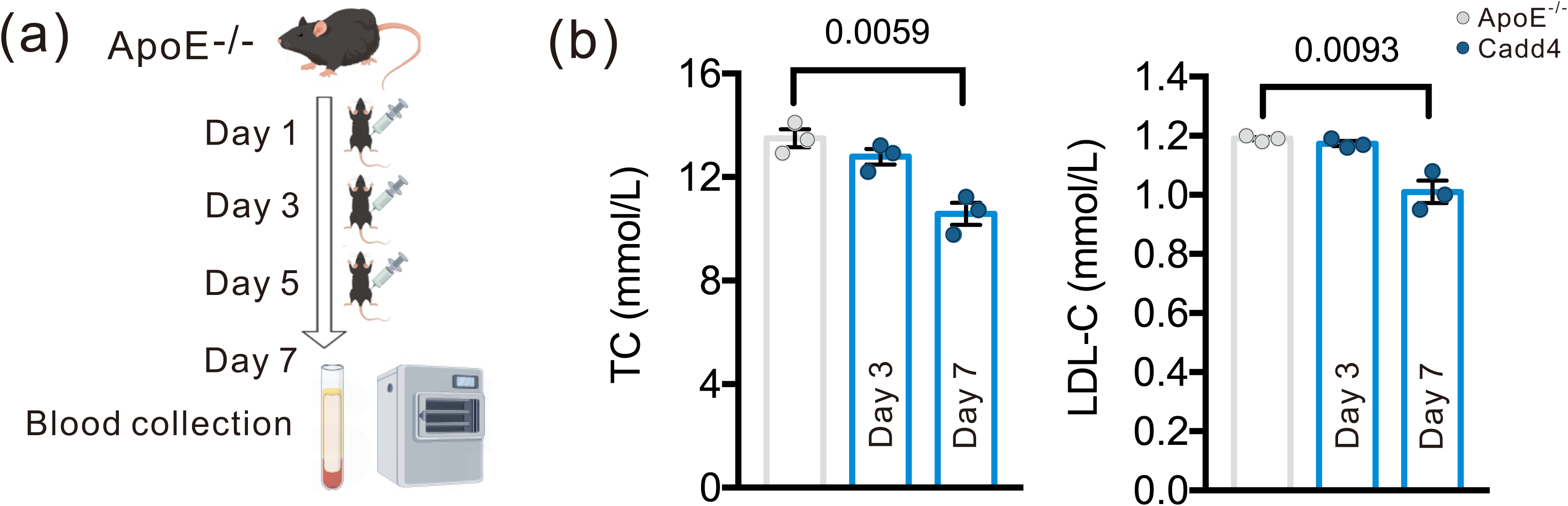
Therapeutic peptide-degrader against plasma cholesterol in ApoE^-/-^ mice. **(a)**, experimental design of Cadd4 administration in ApoE^-/-^ mice is shown. **(b)**, serum levels of TC, LDL-C in the ApoE^-/-^ mice treated with and without PCSK9 degrader Cadd4 (N=3). (Data were presented as means ± SD).

